# Expert-led priorities for a response diversity research agenda in ecology

**DOI:** 10.1101/2025.09.17.676972

**Authors:** Samuel R.P-J. Ross, Ceres Barros, Laura E. Dee, Mike S. Fowler, Owen L. Petchey, Takehiro Sasaki, Hannah J. White, Anna LoPresti

**Affiliations:** Integrative Community Ecology Unit, Okinawa Institute of Science and Technology Graduate University, 1919-1 Tancha, Onna-son, Kunigami-gun, Okinawa, Japan 904-0495; Theoretical Sciences Visiting Program, Okinawa Institute of Science and Technology Graduate University, Onna, Okinawa, 904-0495, Japan; Pacific Forestry Centre, Canadian Forest Service, Natural Resources Canada, Victoria, Canada; Faculty of Forestry, University of British Columbia, Vancouver, British Columbia, Canada; Department of Ecology and Evolutionary Biology, University of Colorado Boulder, Boulder, Colorado, U.S.A.; Department of Biosciences, Swansea University, Singleton Park, Swansea, U.K. SA2 8PP; Department of Evolutionary Biology and Environmental Studies, University of Zurich, Zürich, Switzerland; Graduate School of Environment and Information Sciences, Yokohama National University, Yokohama, Japan; Institute for Multidisciplinary Sciences, Yokohama National University, Yokohama, Japan; School of Life Sciences, Anglia Ruskin University, Cambridge, U.K.

**Keywords:** Response diversity, ecological stability, functional diversity, expert survey, qualitative coding, free-text analysis, research prioritisation, horizon scan

## Abstract

Response diversity aims to capture and explain the variation in ecological responses to environmental change. Response diversity is expected to drive ecological stability since a wider variety of responses to one or more environmental factors should stabilise fluctuations of ecosystem functions. However, uptake of empirical response diversity research has been slow. Here we assess current thinking around response diversity by conducting a targeted expert survey of response diversity researchers. Our survey revealed that one barrier to a unified research agenda on response diversity is the lack of agreement among respondents on the definition of response diversity, and to which dimension(s) of ecological stability response diversity might relate. When asked to select the temporal, spatial, and biological scales at which response diversity may be most relevant for ecological stability, respondents chose a wide range of scales indicating differences in how experts view response diversity’s stabilising effect. Respondents considered studies incorporating both biotic interactions and abiotic environmental responses to be especially challenging, as were those thinking about responses to multiple environmental changes simultaneously. Moreover, respondents thought inconsistencies in the definitions of, and methods for measuring, response diversity were a major challenge facing the field. Despite these barriers, the survey revealed a desire for globally coordinated research efforts on response diversity in the form of syntheses, workshops, and distributed experiments, but that a standardised response diversity metric across diverse use-cases could be too restrictive. Our findings suggest we can shift response diversity from a loose collection of conceptual studies and inconsistent empirical applications towards a quantitative and coordinated research programme mechanistically linking biodiversity and ecological stability. As such, we are launching the Response Diversity Network—a research community interested in the science and application of response diversity— whose activities we hope will benefit both individual studies of response diversity and globally coordinated research efforts.

## Background

Ensuring stable and reliable ecological processes that maintain the natural world and nature’s contributions to people is essential for global sustainability. One potential mechanism underpinning the stability of ecosystem processes, functions, and services in the face of environmental change is response diversity. Response diversity describes variation in ecological responses to environmental conditions (Elmqvist et al. 2003; Mori et al. 2013), with higher response diversity indicating a wider variety of organism-environment responses. Response diversity theory is grounded in the insurance effect of biodiversity, which posits that declines of some organisms in response to an environmental change can be offset by neutral or positive responses of others if these organisms have different environmental tolerances and affinities (Yachi & Loreau 1999). In turn, this response diversity should prevent dramatic losses or extreme fluxes of biomass through such compensatory effects (Mori et al. 2013). Response diversity is thus conceptually linked to various dimensions of ecological stability and sustainability (Elmqvist et al. 2003; Mori et al. 2013; Nyström 2006; Ross & Sasaki 2024; Walker et al. 2023).

Empirical evidence mechanistically linking response diversity to ecological stability is rare. Key studies conceptualise links between response diversity and stability (Mori et al. 2013), yet a small collection of studies has empirically measured response diversity using a variety of methods for quantifying and analysing response diversity (Ross et al. 2023). While theory increasingly recognises that functional traits can underlie environmental responses (de Bello et al. 2021; Oliver et al. 2015; Suding et al. 2008), studies using functional traits or other methods to measure response diversity *per se* are less common. Measuring response diversity typically involves either measuring the diversity of functional response traits or differences in species–environment relationships in the field (Suding et al. 2008; Winfree & Kremen 2009) or quantifying differences in how some aspect of a species’ performance relates to the environment (Leary & Petchey 2009; Ross et al. 2023).

Recently, Ross and Sasaki (2024) suggested that the slow uptake of response diversity in empirical studies is caused by several limitations to the concept’s applicability, including inconsistencies in the definition of response diversity across studies, lack of clear methodologies for measuring response diversity empirically, and different ideas around how response diversity should relate to different dimensions of ecological stability such as temporal variability or resilience in the face of disturbance (see Donohue et al. 2016). Some of these limitations may be due to lack of standardisation, coordination, and communication among researchers working on response diversity, resulting in a patchy and disjointed response diversity literature (Ross & Sasaki 2024). However, whether these views on the current challenges and limitations of the response diversity concept are held by experts in the field more broadly remains an open question.

Here, we aim to explore how researchers already working in the emerging field of response diversity (our target population) understand and study response diversity, and determine to what degree the concept—as currently formulated and understood—is operationalizable for empirical study and conservation and management applications. To do so, we surveyed response diversity experts about definitions and concepts surrounding response diversity, current and ongoing empirical response diversity research, and challenges limiting uptake of response diversity in theoretical and empirical studies. Our results should serve as a roadmap for the future development of response diversity as a framework for predicting ecological stability and with application to conservation and ecosystem management.

### Survey methods and analysis

#### Overview of the expert survey

Given that our target population was researchers already working on response diversity, we surveyed experts identified from the response diversity literature and members of the recently formed Response Diversity Network (https://responsediversitynetwork.github.io/RDN-website/), our study population (Appendix 1). We conducted our survey in August-September 2023 via the Qualtrics platform using a combination of free-text, multiple choice, sliding scale, and ranking survey questions (Appendix 2). We asked respondents how clear and well-defined they thought the response diversity concept is, how they thought it relates to ecological stability concepts and management actions, and about their own work on response diversity. To explore how expert respondents consider response diversity to be affected by scale, we provided a hypothetical case study where respondents chose the temporal, spatial, and biological scales they deemed most relevant to “measure, monitor, or manage for persistence or stability”. Our case study involved the persistence of different plant species, each with their own environmental tolerances (niches), and aggregate temporal stability as a desirable property of the community to ensure stable provisioning of ecosystem functions and services through time (see Appendix 2 for exact wording of the case study). Finally, we asked forward-looking questions to identify the largest challenges facing current studies of response diversity and what (if anything) respondents would like to see as potential outcomes of global collaborative efforts on response diversity.

We recognise the limitations of conducting the literature search and survey in a single language (English), which may reduce respondent cultural and geographic diversity. Given that response diversity is a relatively young field, lack of established terms hinders comparable translation across languages, which can undermine conceptual equivalence (Squires 2009). Moreover, accurate translation for domain-specific scientific terms is a well-known challenge, even within well-established fields of research (Naveen & Trojovský 2024). We therefore conducted our research in a single language but suggest this gap be addressed in future work as the field continues to refine terms across languages and awareness of the concepts grows across different stakeholder groups (see ‘Future Directions’).

#### Qualitative and Quantitative Analyses

To analyse our survey results, we used a mix-methods approach including quantitative analysis of numerical and ordinal (ranked) responses and inductive content analysis—a type of qualitative coding—to identify patterns and themes from open ended (free text) questions (Kyngäs 2020).

Further details of survey design, implementation, analyses of quantitative and qualitative data, and respondent demographics, are presented in Appendix 1. Here we report results from this expert survey, and draw on them to shape efforts of the recently formed Response Diversity Network (https://responsediversitynetwork.github.io/RDN-website/), which aims to coordinate future research based on researchers’ community-driven priorities for response diversity.

#### Respondent demographics and expertise

We received responses from 69 experts across career stages, fields of ecology, and geographic locations (Appendix 1, Supplementary Table S1). Our response rate was 18.3%, likely reflecting the length and complexity of our survey, both factors known to reduce response rates. Nevertheless, given the small size of the response diversity field and that our study population was published experts of response diversity and members of the new Response Diversity Network, our sample size was appropriate for our purpose and is the largest sample of response diversity experts surveyed to date.

Our study population was dominated by researchers from Europe (53.7% of responses) and North America (30.0%), with no responses from researchers in Africa. This is despite having sampled broadly within the response diversity literature (Appendix 1) and initially inviting researchers from across the globe to join the Response Diversity Network. Our response rate was lower for researchers based in the Global South (∼5.3%) than those in the Global North (∼20.6%; binomial GLM with logit link: Coef =-1.54 ± 0.61 SE, *z* =-2.53, p = 0.012), perhaps reflecting the disproportionate time burden placed on historically underrepresented scholars and those working at underfunded institutions (Akin 2020; Heng et al. 2023). Yet, our survey pool, and hence the response diversity literature, is also significantly biased towards researchers from the Global North (exact binomial test, two-sided against *p* = 0.5: *n* = 377, *k* = 316, estimated probability = 0.838 (0.797 – 0.874 95% CI), p < 0.001). We also asked survey respondents to report the study locations of their response diversity research (Question 8; Appendix 2). Of the 71 studies reported, 25 were not location-specific (*e*.*g*., they were theory, experiments, or data syntheses). Of the remaining 46 studies, only 8 studies (17.4%) were conducted primarily in countries in the Global South. Thus, we conclude that the response diversity literature is currently skewed towards authors from the Global North and study systems based in the Global North, which is reflected in the composition of our study population.

Respondents generally identified as more familiar with the concept of ecological stability than with response diversity (Figure 1; Tukey pairwise comparisons between logit transformed means = 1.729, S.E. = 0.438, p < 0.001). In contrast, there was no evidence of a significant difference between respondents’ self-reported familiarity with ecological stability and functional diversity (difference = 0.966, S.E. = 0.438, p = 0.070), nor functional diversity and response diversity (difference = 0.763, S.E. = 0.366, p = 0.094). Respondents’ familiarity rankings for ecological stability and functional diversity were not significantly correlated (Spearman’s rS = 0.005, p = 0.965), nor were the familiarity rankings attributed to ecological stability and response diversity (rS = 0.042, p = 0.732); there was, however, a weak, positive relationship between individual familiarity rankings for functional diversity and response diversity (rS = 0.364, p = 0.002). Respondents, who are experts publishing on the topic of response diversity in the academic literature, reported lower familiarity with response diversity than with the concept of ecological stability, perhaps suggesting that response diversity is still conceptually ill-defined or less widely discussed than stability.

**Figure 1.**
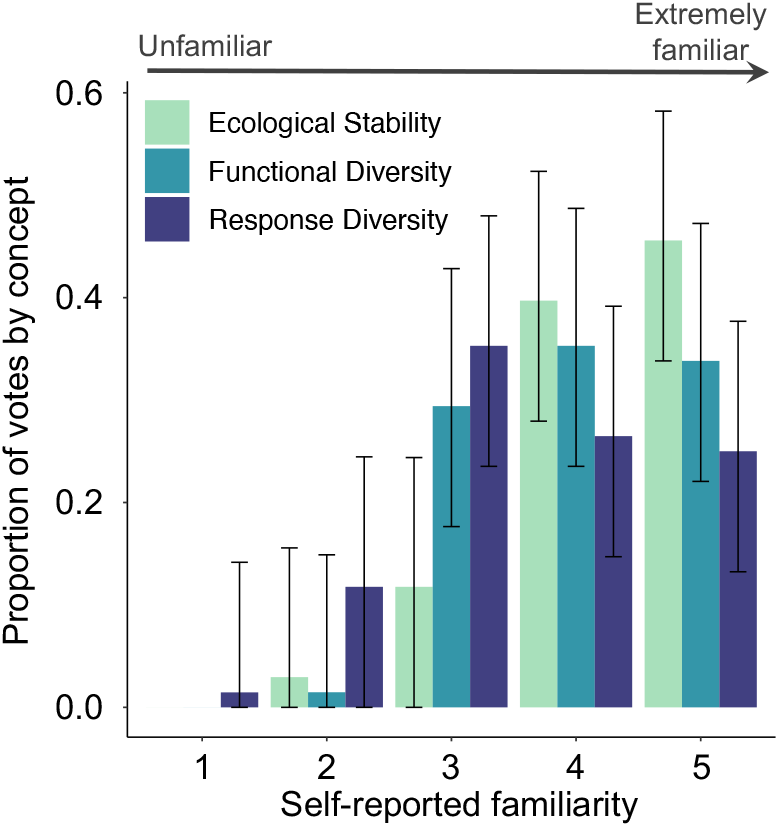
Respondent expertise. Respondents’ self-reported familiarity with response diversity and two related concepts: ecological stability and functional diversity (survey question 1 in Appendix 2). Higher numbers represented greater familiarity with the concepts: 1 = Unfamiliar; 2 = Slightly familiar; 3 = Moderately familiar; 4 = Very familiar; 5 = Extremely familiar/actively work in that field. Y-axis values are the proportion of votes per concept ± 95% multinomial confidence intervals.

### Expert perspectives on response diversity concepts and definitions

#### Conceptual clarity

Here we discuss respondent perceptions related to the concept and definition of response diversity. A concept is the broader, often more abstract idea, whereas a definition is the concise, concrete wording used to clarify that idea. Definitions are tools that help us communicate concepts clearly and consistently. When asked about the clarity of response diversity concepts and definitions, 41% (95% binomial C.I. = 30-52%) of the 69 respondents said the concept was clear, while 45% (C.I. = 34-57%) said it was not. The probability that respondents’ answers to conceptual clarity was either clear, unclear, or unsure was not affected by career stage (multinomial GLM Likelihood ratio c^2^ = 10.636, d.f.= 8, p = 0.223), geographic location (c^2^ = 7.385, d.f. = 8, p = 0.496), or the number of subject areas they aligned with (c^2^ = 0.570, d.f. = 2, p = 0.752).

When asked to provide comments on how the concept of response diversity should be clarified, qualitative coding of open-ended responses revealed that respondents mostly addressed conceptual clarity (48%, 12/25 respondents) and definitional clarity (44%, 11/25), followed by the methodological clarity (36%, 9/25), and clarity regarding the analytical scale of relevance (12%, 3/25). Specifically, those suggesting clarifying the concept did so because response diversity was ambiguous, vague, or missed important considerations such as biotic interactions. Those focusing on the definition said response diversity was unclearly defined, has multiple definitions, or is defined in a way that is not mathematically or empirically tractable (Appendix 3). For instance, 16% (4/25) of respondents used the phrase “response to what?” in their free-response comments, underscoring that response diversity should be defined in relation to some particular environmental variable, change, or disturbance (Elmqvist et al. 2003).

When asked whether response diversity has a widely accepted definition, most respondents said no (55%, 38/69) or were unsure whether a widely accepted definition exists (30%, 21/69). When presented with a choice of definitions and asked to choose any that most closely matched what they consider response diversity to be (Fig. 2), 33% (50/153) of respondents identified response diversity as the “Variation in how *species* respond to environmental change”, 21% (32/153) the “Variation in how *individuals* respond to environmental change”, 20% (31/153) the “Variation in how *functionally redundant species* respond to environmental change”, 18% (27/153) the “Diversity of *functional response traits* within a functional effect group…”, 8% (13/153) the “Variation in *gene expression* under environmental change”, while two respondents did not select any of these options. This skew towards defining response diversity based on species responses likely reflects the participant pool of the survey; most of our respondents self-identified as community ecologists (Appendix 1, Supplementary Table S1) and so would perhaps be most likely to think of response diversity in this context. Alternatively, influential reviews have typically focused on response diversity in the community ecology context (e.g., Elmqvist et al. 2003; Nyström 2006; Mori et al. 2013), which could explain this preference to define response diversity among species responses. Future efforts to define response diversity beyond the community context should prove fruitful.

**Figure 2.**
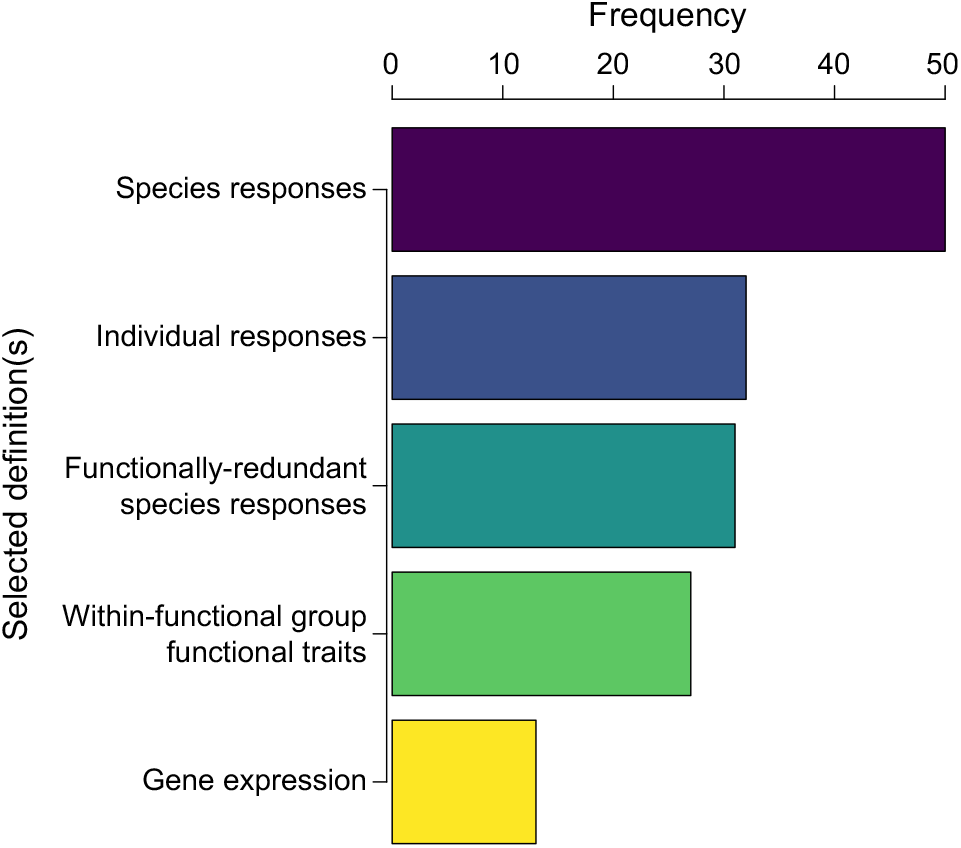
Response diversity definitions. Frequency that survey respondents selected different possible definitions of response diversity (Survey Question 4; Appendix 2). All definitions generally included the idea of variation in responses of some ecological unit to environmental change, with specific differences captured here by the paraphrased options (see Appendix 2 for full definitions). Note that respondents could select multiple definitions. “Other” was selected 10 times and respondents then provided their own response.

Among the remaining 10 respondents who thought response diversity has a widely accepted definition, there was not agreement about what this definition is; respondents mentioned “organisms”, “species”, “communities”, and “functional groups” in their responses equally often.

Several respondents chose or provided response diversity definitions that included the concept of functional grouping; that is, these respondents bounded the definition by necessitating that species be functionally similar, such as “…response diversity among functionally redundant organisms will favour stability”. While older definitions of response diversity in ecology included this need for functional similarity (Elmqvist et al. 2003; Laliberté et al. 2010), subsequent studies defined response diversity without such constraints (Mori et al. 2013; Ross et al. 2023).

One idea that appeared across different responses was that response diversity “must be understood as an emergent feature of a complex system”. This is appealing since systems with more species—or individuals in the case of genetic or intraspecific response diversity (Herrando-Pérez et al. 2019; Kahiluoto et al. 2019)—may be generally expected to have higher response diversity arising from niche differences that reflect environmental tolerances. However, emergence is a feature of complex systems that necessitates a multi-scale approach to its empirical study (de Haan 2006); in the case of response diversity, this means both a focus on the unit of interest (genes, individuals, species) and on higher-level (community) response diversity. Integrating research across scales is challenging; considering response diversity emergent may limit the applicability of the concept for conservation and management.

Respondents seemingly advocated for standardisation of methods for measuring response diversity (50%, 28/56 respondents), while some expressed doubts about whether such standardisation should be pursued (21%, 12/56)—reflecting the philosophical state of the field—or could be practically achieved (13%, 7/56). When asked to identify any standardised methods for measuring response diversity, four respondents pointed to our recent attempt to develop a general framework for measuring response diversity from demographic rates and performance-environment relationships (Ross et al. 2023), and one respondent indicated Darling et al. (2013), who apply the delta method (Oehlert 1992) to estimate the variance of effect sizes for coral cover change with fishing pressure. In open ended questions, concerns arose that standardisation would restrict creative freedom and that it is unlikely a single method would be applicable across different study systems, objectives, and scales (34%, 19/56).

#### Links with stability and management

Ecological stability is itself a multidimensional concept with a plethora of specific metrics used to measure stability across contexts (Donohue et al. 2016). When asked how the response diversity concept should be clarified, six respondents provided free-text responses, of which two mentioned that the underlying stability concepts on which response diversity is built are themselves overly complex or unclear. In support of this, when asked to select which dimension(s) of stability they expect response diversity to relate to, 67 respondents selected from nine potential descriptions 327 times (Figure S2). The frequency of selecting each definition differed from a uniform distribution (c^2^ = 21.117, d.f. = 8, p = 0.007; Figure S2). 68% (47/69 respondents) thought response diversity should relate to the persistence of ecosystem service delivery; 67% (46/69) to community or emergent property persistence; 65% (45/69) to asynchrony of population dynamics among species in a community; 56% (39/69) to reactivity to abrupt change; 53% (37/69) to temporal variability of communities or emergent ecosystem properties; 53% (37/69) to ecological resilience; 41% (28/69) to spatial asynchrony; 39% (27/69) to engineering resilience; and 31% (21/69) to spatial variability or patchiness (Figure S2a). One respondent did not expect response diversity to relate to stability, while 91% (63/69) thought response diversity may mechanistically drive two or more dimensions of ecological stability (Figure S2b). These results suggest a general feeling that response diversity should drive persistence and stability of aggregate ecosystem properties, but indicate less support for response diversity’s role in spatial structure or resilience. We aimed to further disentangle the drivers of respondent choices here, exploring why respondents selected certain stability dimensions as potentially being driven by response diversity. There was no evidence that the chosen dimension(s) of stability were driven by respondents’ choice(s) of response diversity definition (Question 4; Appendix 2), nor by self-reported interest in understanding stability and its drivers (Question 5 option C). There were some differences by respondents’ area(s) of specialism (Question 19), though given the unbalanced and low sample sizes across some categories, these patterns were not clear (see Appendix 1 for details). Additional theoretical and empirical work will reveal the specific dimensions of stability to which response diversity relates and under which conditions.

Moving from theory to practice, respondents were asked to choose how they see response diversity relating to conservation or management (Appendix 2). 59 respondents selected the following descriptions a total of 112 times: “mitigating the impacts of environmental change on ecosystems based on predictions from response diversity theory” was selected 39% (44/112) of the time; “monitoring response diversity as a neglected aspect of biodiversity monitoring” 34% (38/112); and “maintaining ecosystem functioning under environmental change by active manipulation of community composition (or genetic, or intraspecific variation)” 27% of the time (30/112). These frequencies did not differ from a uniform distribution (c^2^ = 3.964, d.f. = 2, p = 0.138). When asked to explain why they chose these answers, 59% (26/44 respondents) believed that response diversity is an inherently useful indicator to understand and possibly manage ecological change and therefore should be included in management. However, 11% (5/44) suggested more research is needed to better understand how response diversity may be useful for management and whether “…by maximizing response diversity we can build more ecologically resilient ecosystems”. In contrast, 14% (6/44) expressed that active manipulation of community composition to maximise response diversity and in turn promote resilience is logistically intractable or inadvisable unless “clear and specific drivers” are expected to impact a community. Respondents worried that without a clear understanding of the strength and direction of environmental change, active manipulation could be “very dangerous [and] could cause ecosystem collapse” because, for example, “responses to drought might differ from responses to fire [or] grazing”.

#### Response diversity across temporal, spatial, and biological scales

We provided respondents with a hypothetical case study where persistence of plant species with different niches and their temporal stability through time might be desirable properties of a community for stable provisioning of ecosystem functions and services (Appendix 2). We asked respondents to choose the temporal, spatial, and biological scales they deemed most relevant to “measure, monitor, or manage for persistence or stability,” and to indicate how confident they were in their answers (following Isbell et al. 2023; Figure 3). We found that the median relevant temporal scale selected corresponded roughly to an annual scale (a score of 64.5, interquartile range 47 – 75), with a confidence of 59.5% (44 – 75%; Figure 3a). There was a weak, positive correlation between the selected temporal scale and confidence scores (rS = 0.275, n = 66, p = 0.025). When asked why respondents chose the scale they did, qualitative coding revealed that 56% (36/64) of respondents made their decision based on the intrinsic characteristics of the ecological community such as “generation times” or “species turnover”, while 20% (13/64) instead chose based on the characteristics of the environment or disturbance. Only 5% (3/64) chose based simultaneously on the characteristics of the community and the environment. Respondents equally often (14%, 9/64) mentioned the need for temporal scales to encompass either environmental stochasticity or evolutionary processes.

**Figure 3.**
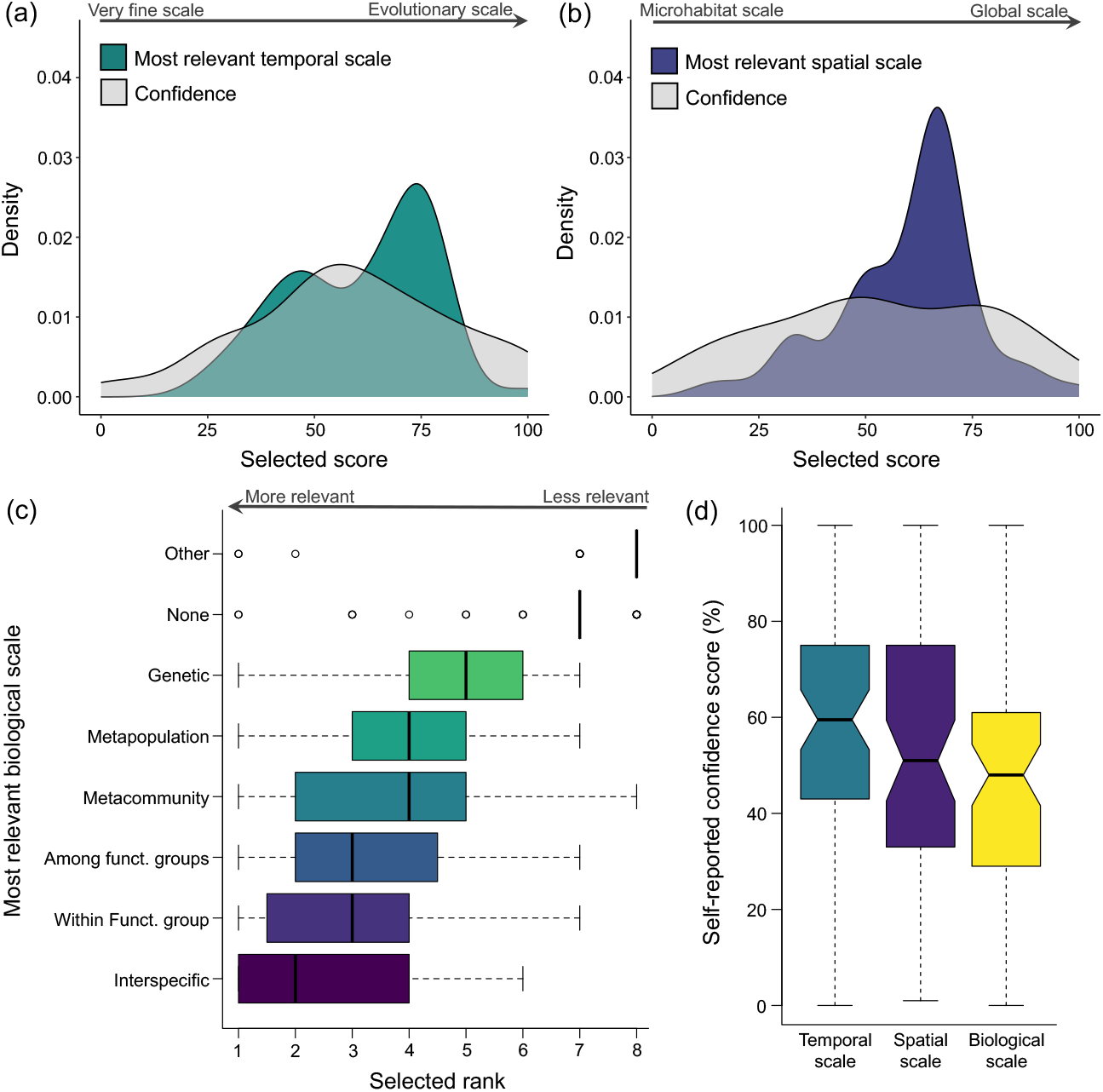
Response diversity and scale. Respondent choices of the most relevant scales to measure, monitor, or manage for persistence or stability in a hypothetical case study (see survey questions 9-11; Appendix 2). Respondents were asked to choose the most appropriate temporal scale (a), spatial scale (b), and biological scale (c) for the case study, as well as to report their confidence level on a scale from 0-100 where 0 is not at all confident, and 100 is extremely confident/certain. Temporal (a) and spatial (b) scores were chosen on a sliding scale from 0-100, where 0 represented very fine temporal (e.g. minutes, hours) or spatial (microhabitats) scales and 100 very large temporal (evolutionary/multiple generations) and spatial (global) scales (see Appendix 2 for details). Panels (a) and (b) are density plots showing the distribution of respondent’s selected scale scores in colour, and selected confidence scores in grey. For biological scale (c), respondents were asked to rank a range of biological scales from most to least relevant for the case study, where 1 indicated most relevant and 8, least relevant. Panel (c) shows the distribution of rank choices per biological scale indicated by box plots with whiskers extending to the most extreme data point which is no more than 1.5 times the interquartile range from the box. Panel (d) compares the distribution of self-reported confidence scores for each dimension of scale as boxplots with the same parameters as in (c).

We found that the median relevant spatial scale selected corresponded roughly to regional or metapopulation scales (median score 66, interquartile range 50 – 68), with median reported confidence of 51% (33 – 75%; Figure 3b). There was no evidence of a correlation between the selected spatial scale and confidence scores (rS = 0.222, n = 62, p = 0.083). As with temporal scale, 60% (34/57) of respondents made their choice based on the characteristics of the community, followed by the characteristics of the environment (28%, 16/57). Respondents more often chose spatial scales based on both the characteristics of the community and the environment (14%, 8/57) compared to when choosing the most relevant temporal scale (5%, 3/64). Similarly, more respondents made their choice based on practical or logistical considerations such as “coordinating actions at [this] scale seems more feasible” when selecting the most relevant spatial (21%, 12/57) compared to temporal scale (13%, 8/64). This may indicate that spatial constraints are more stringent than temporal ones when designing experiments or management plans.

Finally, when asked about biological scale, there were significant differences in the ranks awarded to each biological scale (Kruskal-Wallis c^2^ = 265.32, d.f. = 7, p < 0.001). Dunn’s post-hoc pairwise comparisons (using Benjamin-Hochberg adjusted p-values) revealed interspecific response diversity was consistently ranked higher—considered more relevant to the persistence of functioning in the case study—than all other biological scales, except within- and among-functional groups, and metacommunity scale response diversity (Figure 3c). Here, respondents chose the most relevant biological scale based less on the characteristics of the community (14%, 7/50), and more on the relevance of the biological scale to ecosystem functioning or stability (36%, 18/50). These included points such as, “[for] stable biomass production in a community, interspecific or functional group response diversity might be most relevant”. That respondents mainly focused on interspecific response diversity, functional group response diversity, and metacommunity response diversity may again reflect our respondent pool’s focal research interests (Appendix 1, Table S1). For both spatial and biological scales, respondents emphasised the need to consider multiple scales simultaneously, whereas they did not for temporal scale. This is perhaps because temporally down-sampling a high-resolution time series *a posteriori* to capture different temporal scales does not require specific sampling designs, whereas sampling at multiple scales in space is less straightforward, and sampling at multiple levels of biological organisation demands *a priori* cross-scale design.

We found a significant correlation between survey respondents’ confidence scores corresponding to temporal and spatial scales (rS = 0.565, n = 62, p < 0.001). That is, researchers who were more confident in their choice of temporal scale also tended to be more confident in their choice of spatial scale, but there was no evidence of a correlation between the selected spatial and temporal scale values themselves (rS = 0.240, n = 66, p = 0.052). There was a difference in self-reported confidence across temporal, spatial and biological scales (KW-test c^2^ = 7.67, df = 2, p = 0.02; Figure 3d). Dunn’s multiple comparisons test provided evidence for a significant difference in the self-reported confidence scores for biological and temporal scale values (Z =-2.72, adjusted p = 0.02), but not between confidence in spatial and temporal scale values (Z =-0.86, adj. p = 0.39) or between biological and spatial scale values (Z =-1.82, adj. p = 0.10). Overall, very few respondents had high confidence scores across any of the three dimensions of scale they were asked to consider and respondents broadly selected values from the full range of temporal, spatial, and biological scales.

Together, these results suggest differences in how experts view response diversity’s potentially stabilising role across different spatiotemporal scales and levels of biological organisation.

### Future directions: a roadmap for the Response Diversity Network

Multiple barriers to response diversity research identified by Ross and Sasaki (2024) were also identified by survey respondents, including inconsistencies in the definition of response diversity, in how it should be expected to relate to stability, and in its measurement. Biotic interactions can also complicate the study of response diversity; organismal responses to the environment are shaped not only by the (multiple) environmental axes that affect species, but also by biotic interactions. Realised environmental responses therefore represent an organisms’ inherent environmental response modified by any intra- or interspecific interactions, which in turn also depend on the environment (Fox & Morin 2001). Moreover, the temporal, spatial, and biological scale of focus affects the observation of traits, environmental responses, and ecological stability (Clark et al. 2021; Gonzalez et al. 2020), hindering development of a multi-scale framework for considering response diversity. Together, these factors, as well as others, curtail the applicability of response diversity as a practical concept for ecology and sustainability science (Ross & Sasaki 2024; Walker et al. 2023).

We asked survey respondents to identify the main challenges facing studies of response diversity (Appendix 2). There were significant differences in the ranks awarded to each challenge (Kruskal-Wallis c^2^ = 214.52, d.f. = 7, p < 0.001), with Dunn’s post-hoc pairwise comparisons (using Benjamin-Hochberg adjusted p-values) showing that complexity arising from interspecific interactions was consistently ranked higher than other challenges, except (lack) of clarity around definitions/aims, and multiple environmental stressors (Figure 4a). Recent research is beginning to address these challenges. While Ruiz-Moreno *et al*. (2024) test the relative importance of response diversity and species interactions for driving reef fish dynamics (see also Ives & Carpenter 2007), only one recent study explicitly addresses importance of ecological interactions for response diversity, finding that interspecific interactions confound response diversity’s stabilising effects (Kunze et al. 2025). There are also ongoing efforts to measure response diversity in a multifarious environmental change context (Polazzo et al. 2024), and others aiming to develop response diversity methods for predicting ecosystem service persistence (Genung & Winfree 2024), coexistence outcomes (De Laender et al. 2023), or responses to diverse perturbations (Orr et al. 2024). Ordinal logistic regression showed that the rankings awarded to each future challenge category varied with career stage (interaction term Challenge:Career Stage; Likelihood ratio c^2^ = 57.52, d.f. = 28, p < 0.001). Excluding one individual who did not report career stage, we find a difference in assigned ranks according to career stage in the “Complexity from ecological interactions” challenge, where mid/late career researchers (n = 31, marginal mean odds log ratio of rank =-46.9, S.E. = 0.445) tend to rank this Challenge more highly than early career researchers (n = 10, marginal mean odds log ratio of rank =-48.3, S.E. = 0.444) (LR c^2^ = 10.016, d.f. = 4, p = 0.040; mean difference in ordinal logs-ratios =-1.437, S.E. = 0.545, z =-2.639, p = 0.042 based on Tukey adjustment for multiple comparisons). No other Challenges differed in the ranks assigned by researchers at different career stages.

**Figure 4.**
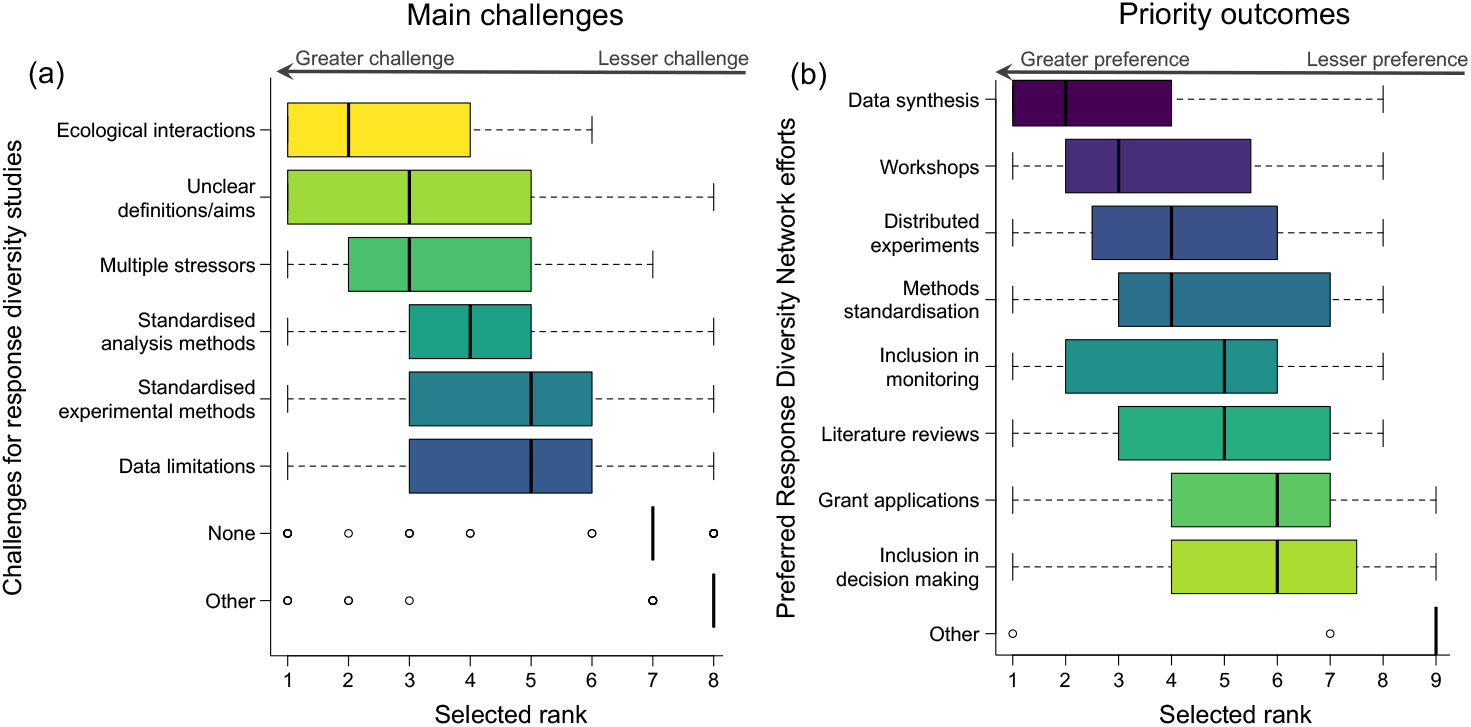
Outlook for coordinated efforts. Respondents’ selected importance rankings of challenges for studies of response diversity (a) and preferred outcomes of global collaborative efforts to coordinate response diversity research through the Response Diversity network. Options are paraphrased here for ease of visualisation, but see survey questions 13 and 16 (Appendix 2) for full descriptions of main challenges and priority outcomes, respectively. Respondents were asked to rank the importance of each challenge (a) or outcome (b), where 1 indicated most important or preferred. The distribution of rank choices per selection is indicated by boxplots, with whiskers extending to the most extreme data point which is no more than 1.5 times the interquartile range from the box.

Next, we introduce the Response Diversity Network (https://responsediversitynetwork.github.io/RDN-website/): a group aiming to stimulate and coordinate research on response diversity and ecological stability. The network is still in its infancy (111 members from 76 institutions in 27 countries at time of writing), so we will use the outcome of this survey to establish core aims and objectives and inform the strategic direction of the Response Diversity Network, and ultimately to address the challenges identified here. To this end, we asked respondents to rank their preferred outcomes of any future global collaborative efforts to advance the science of response diversity in ecology (Figure 4b). Respondents ranked meta-analyses or other data syntheses more highly as a desired Network outcome than all options other than workshops (Krukal-Wallis c^2^ = 210.92, d.f. = 8, p < 0.001, Figure 4b; see Table S2 in Appendix 1 for Dunn’s pairwise comparisons).

Our literature search and survey responses indicate a probable geographic bias in the response diversity field, with an underrepresentation of research from the Global South. Such an imbalance encumbers scientific progress in the field (Nuñez et al. 2021) and can limit the scope and applicability of response diversity concepts and metrics if they remain untested in study systems from many parts of the globe. One key outcome of this study is informing the strategic direction of the Response Diversity Network, which has the potential to address some of the imbalances in global response diversity research. Our study points to response diversity as a small and geographically biased field relative to the wider research community. This may, in part, be due to researchers not explicitly or knowingly working on “response diversity” owing to lack of clarity around concepts and definitions, as our survey results point to. As such, we aim for the Response Diversity Network to address these issues by pursuing the challenges and research foci identified here; for example, by engaging with diverse researchers, practitioners, and stakeholders, such as those from underrepresented groups and geographies. The Network can facilitate and co-ordinate working groups and proposals aiming to fund response diversity research and meetings in the Global South, and the ongoing members’ seminar series—which already accounts for diverse time zones—provides a space for scientific discussion and networking free from the financial burden of attending scientific conferences. This paper therefore serves as an open invitation for anyone interested in the science or practice of response diversity to join the Network; we especially welcome members from underrepresented backgrounds and geographies, including to serve on the Network’s steering committee and help shape the strategic direction of the Network.

The challenges and priority research foci identified here included issues with plurality in the definition of response diversity and a lack of standardised methods. Plurality in the definition of response diversity and methods for its quantification represents a challenge to the field through a lack of clarity, confusion, and disagreement. An overly broad definition can be criticised for not meaning anything; too narrow and it excludes diverse, relevant perspectives. One solution is to develop a taxonomy of definitions and descriptions of response diversity (Figure 5). Developing such a taxonomy is needed to ensure response diversity is meaningful, measurable, and broadly applicable, but its success hinges upon widespread uptake and careful application (Grimm & Wissel 1997; Herrando-Pérez et al. 2012). For example, during our survey, respondents clarified response diversity through a focus on biological scale (organisms, species, functional groups, *etc*.) and pointed to the difference between response diversity to a specific environmental driver (“response to what?”) *versus* the total response capacity of a community to any set of future environments (Polazzo et al. 2023). This is a useful distinction because, across open-ended questions, respondents converged on the idea that response diversity to one environmental change axis is not expected to provide resilience against a different environmental axis (Mori et al. 2013), while also suggesting that response diversity is the ability “to respond to all kinds of changed [environmental] conditions.” We suggest this distinction between response diversity and response capacity (Polazzo et al. 2024) is a useful boundary for addressing plurality in definitions and concepts.

**Figure 5.**
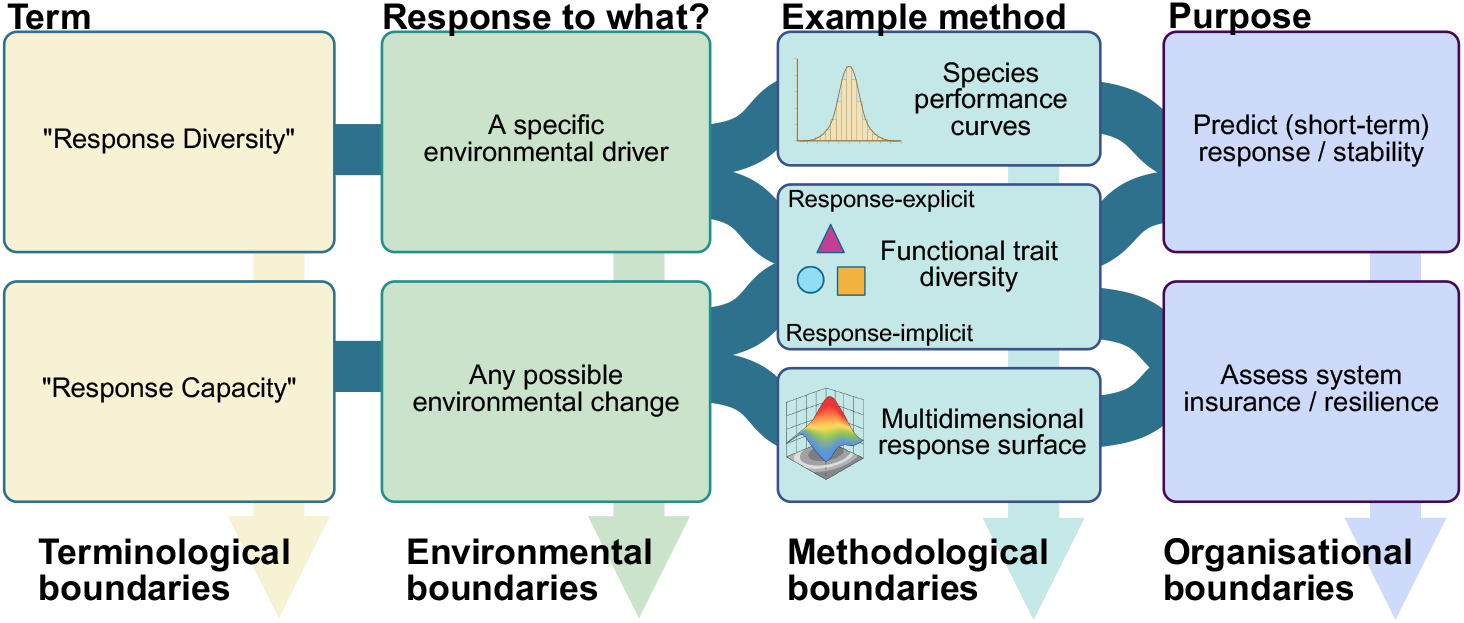
Example taxonomy of response diversity concepts, definitions, and metrics. We suggest adopting a taxonomy to define conceptual and practical boundaries, allowing pluralism and applicability of response diversity concepts while avoiding confusion of definitions and redundant metrics. Terminological boundaries can distinguish similar terms. Environmental (or extrinsic) boundaries provide an answer to the question “response to what?” since response diversity is defined relative to some set of (abiotic or biotic) environmental parameters. Methodological boundaries can be used to separate different methods for measuring response diversity, such as those based on performance curves or those based on functional traits. Under this umbrella, there are also various metrics to measure response diversity; for example, the choice of functional diversity metric (FDis or RaoQ) when using trait-based response diversity methods.

Finally, organisational boundaries include those setting the level of biological organisation of interest (from genetic/molecular/subcellular response diversity to macro-scale), as well as organising the conceptual aims of a given response diversity approach. Figure created with BioRender.

Another example of a clear boundary that could represent a split in the taxonomy is that response diversity measures can be classified based on the data from which they are derived (Ross et al. 2023), including trait data, performance curve data, experimental data, or observational data.

Through open ended questions, respondents were concerned that standardisation of response diversity methods would restrict creative freedom and inhibit application of methods across study systems, objectives, and scales. One solution to this challenge could be “a set of different standardized methods adapted to the different broad types of [response diversity] studies” (Figure 5, ‘Methodological Boundaries’). We suggest this as a key future direction for concerted efforts to standardise response diversity methods and metrics. While it may be unrealistic and/or undesirable to standardise methods across all response diversity research, standardised approaches to using each type of data could increase opportunities for cross-study comparison.

Methods development should focus separately on response diversity metrics for defined classes of use-cases. For example, a first target area for methods development could be evidence-based trait diversity metrics using traits that directly reflect organismal environmental tolerances (Oliver et al. 2015; Suding et al. 2008). A second might be to develop methods based on organismal performance or demographic rates to measure variation among performance-environment relationships (Ross et al. 2023), or to extend this idea across all possible environmental conditions to capture response capacity in a conceptually distinct but complementary methodological framework (Polazzo et al. 2023). Third, given that responses of organisms to environmental conditions should ultimately be driven by their niches, niche-based methods considering variation in niche breadth or optima along environmental axes (*e*.*g*., critical thermal bounds CTmin and CTmax) could produce response diversity methods rooted in environmental niche axes (Fowler & Ruokolainen 2013; Herrando-Pérez et al. 2019). A final suggested avenue is to develop response diversity methods for application to ecological networks; understanding how network properties influence various dimensions of stability remains a key challenge (Domínguez-García et al. 2019; Keyes et al. 2024; Ross et al. 2021). Integrating response diversity into network studies should therefore prove fruitful (Danet et al. 2024).

Here we draw parallels with the literature on ecological stability. Ecologists regularly adapt seemingly vague or nebulous ecological concepts to their specific study systems, in the process producing new methods. For example, from seminal papers on ecological stability (e.g. Holling 1973; Pimm 1984), a suite of possible methodological frameworks and specific stability metrics have arisen to capture a diversity of stability dimensions and closely related concepts (Domínguez-García et al. 2019; Donohue et al. 2016; Grimm & Wissel 1997). It seems respondents want the same when studying response diversity; to adapt the concept of response diversity to their study system using the most suitable metric for each use-case. Accordingly, we suggest one priority is to develop a taxonomy of response diversity concepts. Figure 5 represents one possible starting point for such a taxonomy based on the outcomes of our expert survey. Future research directions also include expanding upon this survey to consider how response diversity terms compare across languages. Such research would contribute to developing cohesion and coordination across the global response diversity community as the Response Diversity Network and possible taxonomies for response diversity concepts and definitions continue to evolve.

Critically, response diversity methods should be developed by grounding them in theory while ensuring they be empirically tractable; collaboration between theoretical and empirical ecology will be key, as will discussion and co-production with practitioners to ensure metrics are useful for monitoring and management. Through efforts to facilitate connections around the world via meetings, funding opportunities, and collaborative projects, and those to establish pathways to include response diversity in observation, monitoring, and policy for sustainability (Walker et al. 2023), the Response Diversity Network aims to both coordinate collaborative work and facilitate independent research, including the development of standardised but flexible methodologies. The Response Diversity Network is growing, and membership is open to anyone interested in ecological stability, its drivers, or related topics. Response diversity is not a new concept (Elmqvist et al. 2003; Nyström 2006) but is gaining more attention in ecology lately, and diverse input is required. Knowledge exchange among the diverse membership of the Response Diversity Network in terms of career stage, research area, and geographic location should help to overcome the breadth of challenges identified in our survey.

As such, we aim for the expert perspectives presented here to set a community driven research agenda for the science of response diversity in future.

## Author Contributions (CRediT taxonomy)

<u>Samuel R.P-J. Ross</u>: Conceptualisation; Data curation; Formal analysis; Investigation; Methodology; Project administration; Validation; Visualisation; Writing – Original draft. <u>Ceres Barros</u>: Conceptualisation; Formal analysis; Investigation; Methodology; Software; Visualisation; Writing – review & editing. <u>Laura E. Dee</u>: Conceptualisation; Investigation; Methodology; Writing – review & editing. <u>Mike S. Fowler</u>: Conceptualisation; Formal analysis; Investigation; Methodology; Software; Visualisation; Writing – Original draft. <u>Owen L. Petchey</u>: Conceptualisation; Data curation; Methodology; Writing – review & editing. <u>Takehiro Sasaki</u>: Conceptualisation; Data curation;

Methodology; Writing – review & editing. <u>Hannah J. White</u>: Conceptualisation; Data curation; Methodology; Project administration; Writing – review & editing. <u>Anna LoPresti</u>: Formal analysis; Investigation; Methodology; Writing – review & editing.

## Statement of Ethics

Our expert survey was approved by Anglia Ruskin University’s School Research Ethics Panel (application number ETH2223-10179) and met all ethical guidelines of all partner institutions, including the review board of the Okinawa Institute of Science and Technology Graduate University (OIST) where personal data was processed and anonymised before analysis. We will not make any identifying information available as part of this article or associated data; only fully anonymised data is available.

## Acknowledgements

We firstly thank 69 anonymous survey respondents for their contributions to this project. The work described in this paper in part results from the activities and support of the Response Diversity Network (https://responsediversitynetwork.github.io/RDN-website/). We also thank Izabela Mihai for advice on survey design and ethics, and Tad Dallas for ideas about formal analysis, and two anonymous reviewers for constructive comments. SRP-JR was supported by subsidy funding to the Okinawa Institute of Science and Technology Graduate University (OIST) and a British Ecological Society Large Grant (LRB22/1007). This research was conducted while C.B., L.D., M.F., and O.P. were visiting the Okinawa Institute of Science and Technology (OIST) Graduate University through the Theoretical Sciences Visiting Program (TSVP). Authors C.B. to H.W. are listed alphabetically.

