## Supplementary material for "Expert-led priorities for a response diversity research agenda in ecology": _

### Appendix 1. Supplementary Methods and results

#### Survey design

To assess the current state of thinking on response diversity, we designed an expert survey titled "Gauging research interest in and perceptions surrounding response diversity". The full list of survey questions can be found in Appendix 2. The survey had three main sections. The first section focused primarily on definitions of response diversity, experts' interest in, and familiarity with, response diversity, and how it might relate to ecological stability. The second section focused on specific studies of response diversity by asking experts to highlight up to five of their own papers or preprints about response diversity, or to share details about work in progress where unpublished. Specifically, if unpublished, we asked respondents for information about focal ecosystem type, taxa of interest, main objective of the studies, trophic complexity, and the temporal and spatial scope of the studies. Section three posed a hypothetical case study concerning a plant community which ecosystem managers hope to manage for continued functioning and service provision. We asked respondents to choose a temporal, spatial, and biological (genes to metacommunities) scale at which respondents felt response diversity might be most relevant for measuring, monitoring, or managing for the persistence or stability of ecosystem functioning in the case study. Following Isbell et al. (2023), we also asked respondents how confident they were in the answers they gave to these questions about scale. Additionally, section three asked respondents to rank the main challenges they see for response diversity studies, as well as seeking expert opinions on whether methods need to be standardised across studies, and finally what, if anything, should be an outcome of any global collaborative efforts on response diversity. The aim of this final question was to inform the strategic direction of the Response Diversity Network in future. Finally, we asked for demographic information on career stage, geographic location, and (sub)field of expertise, which was coarse grain to not endanger the anonymity of respondents. Though we recognise the limitations of surveying experts in only one language (Nuñez & Amano 2021), we conducted and distributed our survey in English, following a monolingual search for the corresponding authors of relevant papers in Web of Science (see below), which we acknowledge as a limitation of our study. That said, a key finding of our study is that there are not widely agreed upon definitions and terminology surrounding response diversity concepts. As such, translating our survey to non-English languages was not tractable.

#### Survey implementation

We defined response diversity experts as corresponding authors of Scopus-indexed academic journal articles either focusing on, or in some way discussing, response diversity, as well as researchers who opted into membership of the recently formed Response Diversity Network at the time of the survey. To identify relevant papers, we conducted a Web of Science search following the same workflow as (Ross et al. 2023). Briefly, we searched Web of Science for “response diversity” or “response variation” in all fields within the “Ecology” category, and supplemented the results with additional papers we knew to be relevant but that were not included in the initial list. We also searched bioRxiv and EcoEvoRxiv preprint servers for “response diversity”. This resulted in a total of 219 relevant articles, from which author S.R.P-J.R. extracted as many author email addresses as possible from the Web of Science listings (noting that some journals allow multiple corresponding authors, while others list all author email addresses). Where unavailable, the email addresses of corresponding authors were sourced directly from the article webpage or preprint server. In combination with the membership of the Response Diversity Network, this workflow produced 429 unique email addresses (377 discounting pingbacks) to which we distributed the survey. Authors of this manuscript did not respond to the survey.

The survey was distributed online using the Qualtrics survey software (www.qualtrics.com), because of its wide variety of logic options including skipping and looping questions, and its intuitive interface on both computers and handheld devices. Invitation emails were distributed automatically via Qualtrics on 10 August 2023. The survey was open to respondents for 32 days, and we received 69 responses between 10 August and 11 September 2023. We staggered the time of day at which emails were sent to maximise the likelihood that experts in different time zones would see them.

#### Analysis of numeric responses

Statistical analyses were carried out in R software v. 4.4.0 (R Core Team, 2024). Plots were created using base R and ‘ggplot2’ (Wickham 2016), with colourblind-friendly colour maps taken from the ‘viridis’ package (Garnier et al 2024). We assessed whether respondents showed any differences in familiarity with the concepts of Ecological Stability, Functional Diversity or Response Diversity with a mixed-effects binomial GLMM using the ‘lme4’ package (Bates et al 2015). Each of these concepts was a level in the predictor variable, the probability of respondents ranking their familiarity as 4 (very) or 5

(extremely familiar) was the binomial response variable and respondent ID was treated as a random effect (intercept). We generated Likelihood ratio $c^2$ test statistics, d.f. and corresponding p-values for the GLMM with the Anova function in the 'car' package and performed Tukey's pairwise comparisons on logit-transformed Concept level means using the 'emmeans' package (Lenth, 2024). We evaluated associations between individual respondents' familiarity rankings in the above three concepts, using Spearman's Rank Correlation. Multinomial 95% confidence intervals used in Fig. 1 were generated with the MultinomCI function in the 'DescTools' package (Signorell, 2024).
Descriptive statistics on the clarity of response diversity concepts and definitions – means and 95% binomial confidence intervals – were generated using the binoconf function in the 'Hmisc' package (Harrell, 2024) and the impact of career stage was used as a predictor variable on the respondents answers to conceptual clarity (clear, unclear or unsure) as the response variable in a multinomial GLM using the 'nnet' package (Venables & Ripley, 2002). We used Pearson's $c^2$ tests to determine whether the frequency of selecting each of nine potential Response Diversity definitions, and three definitions related to conservation or management, differed from a uniform distribution.

We used Spearman's Rank correlations to evaluate associations among respondents' responses to the relevant temporal and spatial scales that apply to Response Diversity, and their corresponding confidence scores for each category. We used Kruskal-Wallis tests to determine whether there were significant differences among the ranks awarded to each of the seven options proposed for the most relevant biological scales, each of the eight options for the main perceived challenges for Response Diversity studies, and each of the nine options for preferred Response Diversity Network efforts. We used Dunn's test of multiple comparisons using rank sums for post-hoc pairwise comparisons in each case (Ogle et al. 2023; Dinno, 2024). Finally, we used ordinal logistic regression to assess whether the rankings awarded to each future challenge category varied with career stage, with the polr function in the 'MASS' package (Venables & Ripley, 2002) and emmeans with a Tukey correction (Lenth, 2024) to assess differences in pairwise comparisons of ordinal log-ratios of ranks across each challenge category. One respondent was excluded from this analysis for failing to include a career stage in their responses.

#### Analysis of free text responses

To capture dimensions of respondent knowledge beyond the scope of numerical and close ended questions, we included open ended questions (OEQs) in the survey. The OEQs generally focused on respondent opinions regarding: 1) the clarity of response diversity concepts and definitions; 2) the temporal, spatial, and biological scales most relevant to a hypothetical response diversity case study; and 3) the main challenges facing studies of response diversity (see Appendix 2).

OEQs complement numerical data by providing rich contextual detail in respondents' own words (Ferrario & Stantcheva 2022; Singer & Couper 2017). We analysed OEQs using qualitative coding, a process of categorizing qualitative data to identify patterns and themes (Vaughn & Turner 2016; Williams & Moser 2019). We applied an inductive content analysis, a ground-up approach in which categories emerge from iterative readings of the responses rather than setting pre-determined categories based on a theoretical framework (Elo & Kyngäs 2008; Kyngäs 2020). We deemed inductive content analysis most appropriate for this study due to the exploratory nature of the topic, which requires flexibility in the identification and categorization of emergent themes (Kyngäs 2020).

We developed unique coding structures for each OEQ, with categories discussed and iteratively refined by two authors (S.R.P-J.R. and A.L.). To reduce bias and increase the validity of analyses, these authors independently reviewed each OEQ response and coded information based on the agreed-upon structure. Consensus was reached on all disagreements through discussion, and the coding structure was adjusted as needed for conceptual clarity. The coding structure for each OEQ is provided as Appendix 3.

**Respondent demographics**

Our survey received 69 complete responses from experts representing diverse career stages, (sub)fields of ecology, and geographic locations. We received a fairly even distribution of responses by career stage: students or recent graduates within 2 years of receiving their PhD made up 10 of our respondents (14%); 8 respondents (12%) were 2-5 years post-PhD (or had equivalent non-academic experience); 19 (28%) were 5-10 years post-PhD or equivalent; while the largest number of respondents (45%, n = 31 respondents) represented the unbounded category for those who were 10+ years post-PhD or equivalent. We asked respondents to select from seven possible research (sub)fields to which they considered their expertise most relevant. Respondents were able to select multiple categories or could choose not to select any. We found that most respondents by far represented expertise in community ecology (72%, n = 50), followed by ecosystem ecology (38%, n = 26), theoretical or mathematical ecology (26%, n = 18), population ecology (22%, n = 15), biogeography or macroecology (19%, n = 13), socioecology or social sciences (6%, n = 4), and lastly evolutionary biology (4%, n = 3). Our expert respondents were primarily geographically located in Europe (52%, n = 36) and North America (29%, n = 20), followed by Asia (10%, n = 7), South America (3%, n = 2), and Oceania (3%, n = 2). For analysis of respondent expertise, see main text.

**Table S1.** Demographic characteristics of Questionnaire respondents. Respondents could choose multiple subject areas (see Figure S1).

| | | | | | | | |
|---|---|---|---|---|---|---|---|
| Location | Asia<br>10.4% (n=7) | Europe<br>53.7% (n=36) | N America<br>30.0% (n=20) | Oceania<br>3.0% (n=2) | S America<br>3.0% (n=2) | | |
| Career Stage (NA = 1) | Current/recent PhD student<br>14.7% (n=10) | 2-5 years post PhD<br>11.8% (n=8) | 5-10 years post PhD<br>27.9% (n=19) | >10 years post PhD<br>45.6% (n=31) | | | |
| Subject area (NA = 1) | Biogeog.<br>18.8% (n=13) | Comm. Ecology<br>72.5% (n=50) | Evol. Biology<br>4.3% (n=3) | Ecosyst. Ecology<br>37.7% (n=26) | Popul. Ecology<br>21.7% (n=15) | Socioeco/ Soc-sci<br>5.8% (n=4) | Theor./ math. Ecology<br>26.1% (n=18) |

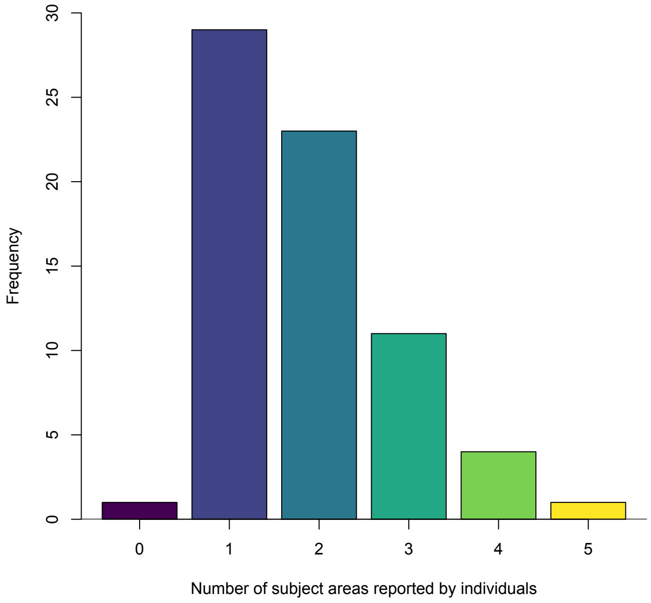


**Figure S1.** How often individuals reported different numbers of subject areas.

### Relationship with different dimensions of ecological stability

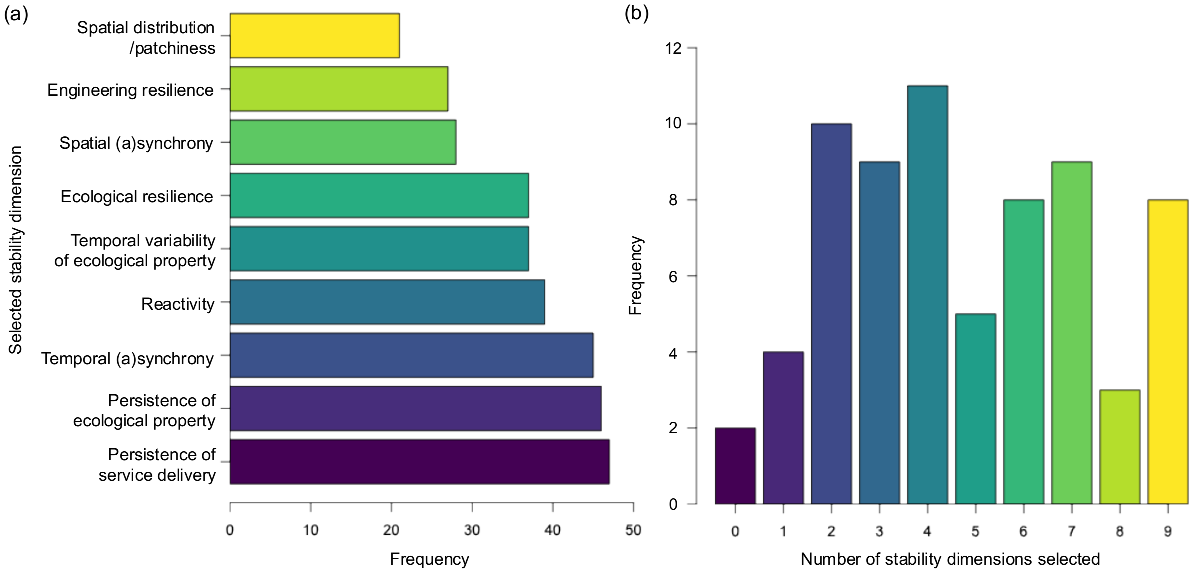


**Figure S2.** (a) Frequency of respondent selections for different dimensions of stability when asked which aspects of stability (if any) response diversity may mechanistically drive (survey question 7; Appendix 2). "None" was selected once, and one respondent did not choose any option (including "None"). Note that respondents could choose multiple stability dimensions; panel (b) shows how many dimensions of stability respondents were choosing. Generally, respondents chose multiple dimensions of ecological stability. Descriptions of stability dimensions are paraphrased for brevity and ease of visualisation, see Appendix 2 for full descriptions.

We tested whether the choice of stability dimension(s) respondents thought response diversity may mechanistically drive (survey question 7; Appendix 2) was determined by three possible drivers: 1) respondents' selected definition(s) of response diversity (survey question 4); 2) whether respondents reported an interest in understanding stability and its drivers as a motivation behind an interest in response diversity (question 5 option C); and 3) respondents' area(s) of research expertise or specialism. Responses to question 5 were converted to a binomial variable coding for whether the respondent did (1) or did not (0) choose option C. One respondent made no selection for question 7 and was excluded. Two others made no selection for question 4 and were excluded from the analysis of the first driver.

We used two complementary approaches to analyse whether the answers to two multiple response questions (i.e., question where respondents can select more than one option, as are questions 4 and 7, and self-reported areas of expertise) were non-randomly related. First Order Rao-Scott Corrected Chi-square tests (available in the svychisq function of the 'survey' package in R) were applied to contingency tables of the two questions to account for the fact that the marginal sums could be greater than the total number of respondents (Decady & Thomas 2000). We also applied separate multinomial GLMs to test each driver's effects (i.e., frequencies of question 4 options, the binary variable of option 5C and respondents' area of expertise) on the probability of choosing either option for question 7 (the response variable). Using the emmeans function in R (Lenth 2024), we then computed and plotted the estimated marginal mean probabilities and their standard errors for pairwise combinations of response and predictor levels. We found no evidence that the selected dimension(s) of stability respondents proposed as possibly being driven by response diversity, were themselves driven by respondents' choice(s) of response diversity definition (adjusted $\chi^2_{36}$ = 5.74, P = 0.44; Fig. S3), nor by self-reported interest in understanding stability and its drivers (adjusted $\chi^2_{9}$ = 1.56, P = 0.97; Fig. S4). We found some differences based on respondents' area(s) of specialism (adjusted $\chi^2_{63}$ = 674.56, P < 0.01; Fig. S5). However, owing to the unbalanced and low sample sizes across respondent area of expertise categories, these patterns were not clearcut (Fig. S5). It is possible that the results were driven by only one respondent selecting option 7A ("No relationship with ecological stability") with that respondent being the only one who selected "Other" in area of expertise."

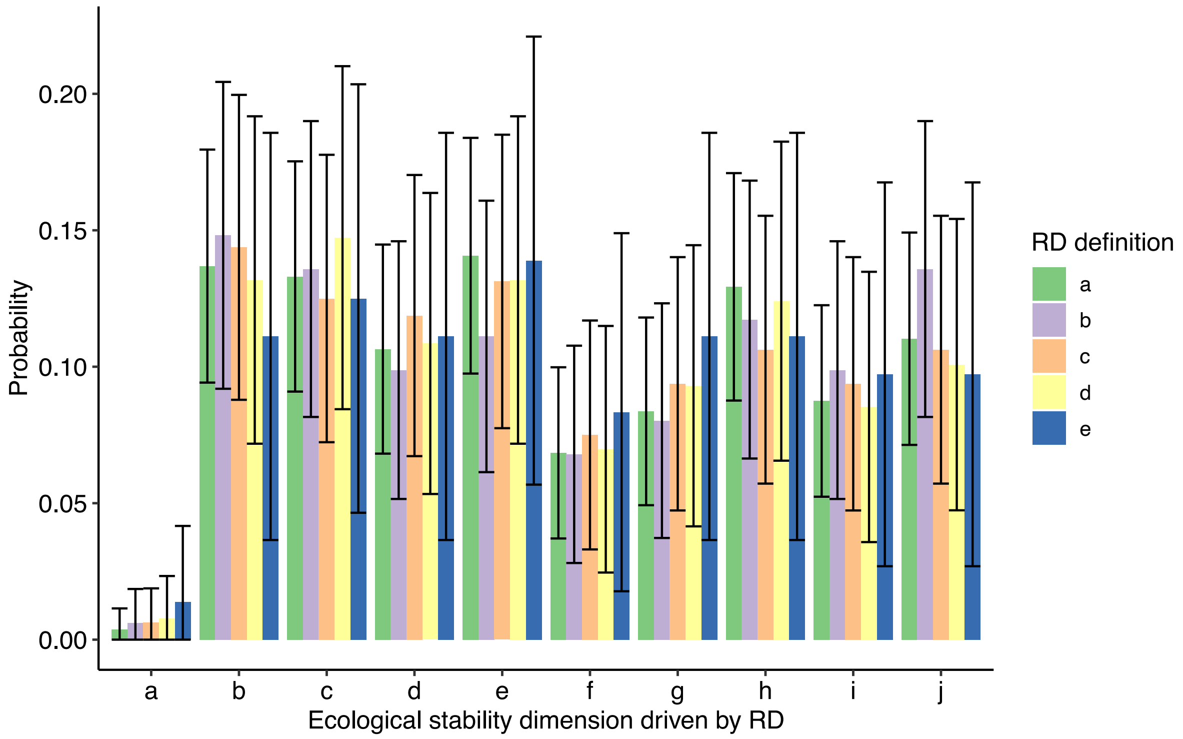


**Fig. S3.** Estimated marginal mean probabilities (bars) and 95% multinomial confidence intervals (error bars) of pairwise combinations of choice of dimension(s) of stability driven by response diversity, RD (question 7, x-axis) and choice of definition of RD in an ecological context (question 4, bar colours). Ecological stability dimensions follow Question 7 (Appendix 2), such that a = no relationship with any stability dimension; b =

persistence of ecological property; c = persistence of service delivery; d = temporal variability of ecological property; e = temporal (a)synchrony; f = spatial distribution/patchiness; g = spatial (a)synchrony; h = reactivity; i = engineering resilience; and j = ecological resilience. Definitions of response diversity follow Question 4 (Appendix 2), such that a = variation in how species respond to environmental change; b = variation in how functionally redundant species (i.e. species contributing to the same ecosystem function(s)) respond to environmental change; c = variation in how individuals respond to environmental change; d = diversity of functional response traits within a functional effect group (a group of species with similar functional effect traits); and e = variation in gene expression under environmental change.

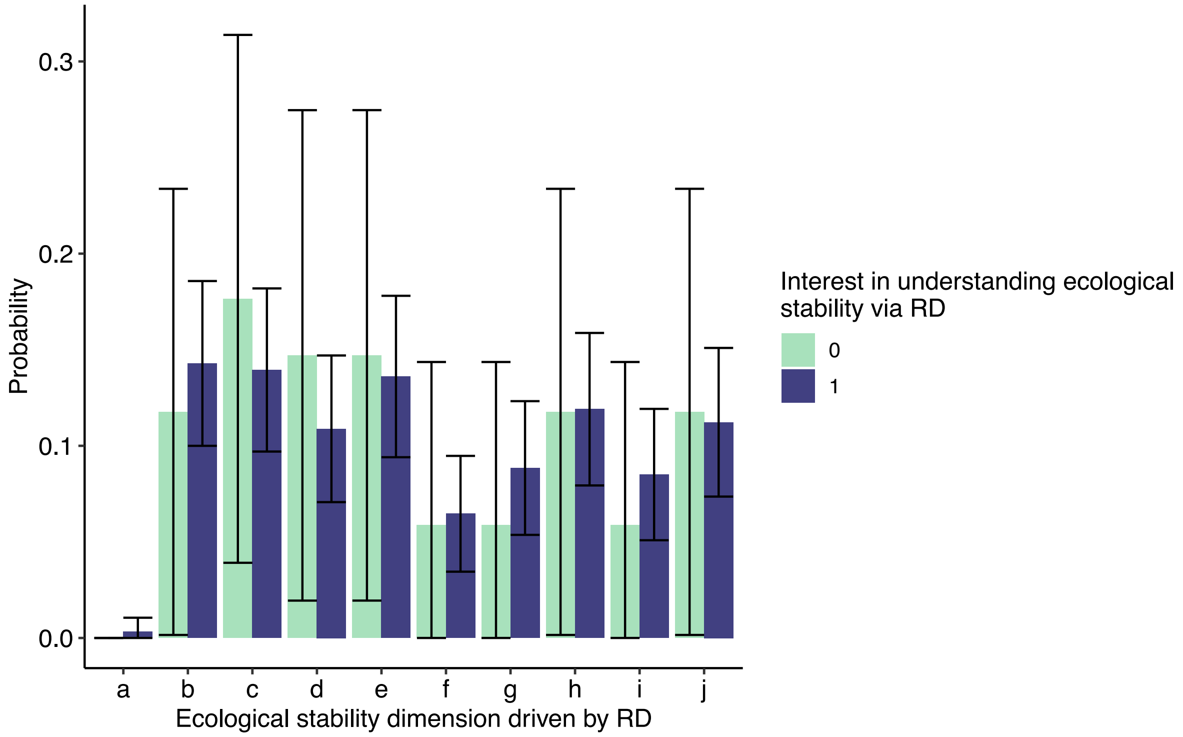


**Fig. S4.** Estimated marginal mean probabilities (bars) and 95% binomial confidence intervals (error bars) of choosing a dimension(s) of stability driven by response diversity, RD (question 7, x-axis) and having/not having an interest in studying ecological stability and its driver using response diversity, RD (question 5 option C, bar colours). Ecological stability dimensions follow Question 7 (Appendix 2), such that a = no relationship with any stability dimension; b = persistence of ecological property; c = persistence of service delivery; d = temporal variability of ecological property; e = temporal (a)synchrony; f = spatial distribution/patchiness; g = spatial (a)synchrony; h = reactivity; i = engineering resilience; and j = ecological resilience.

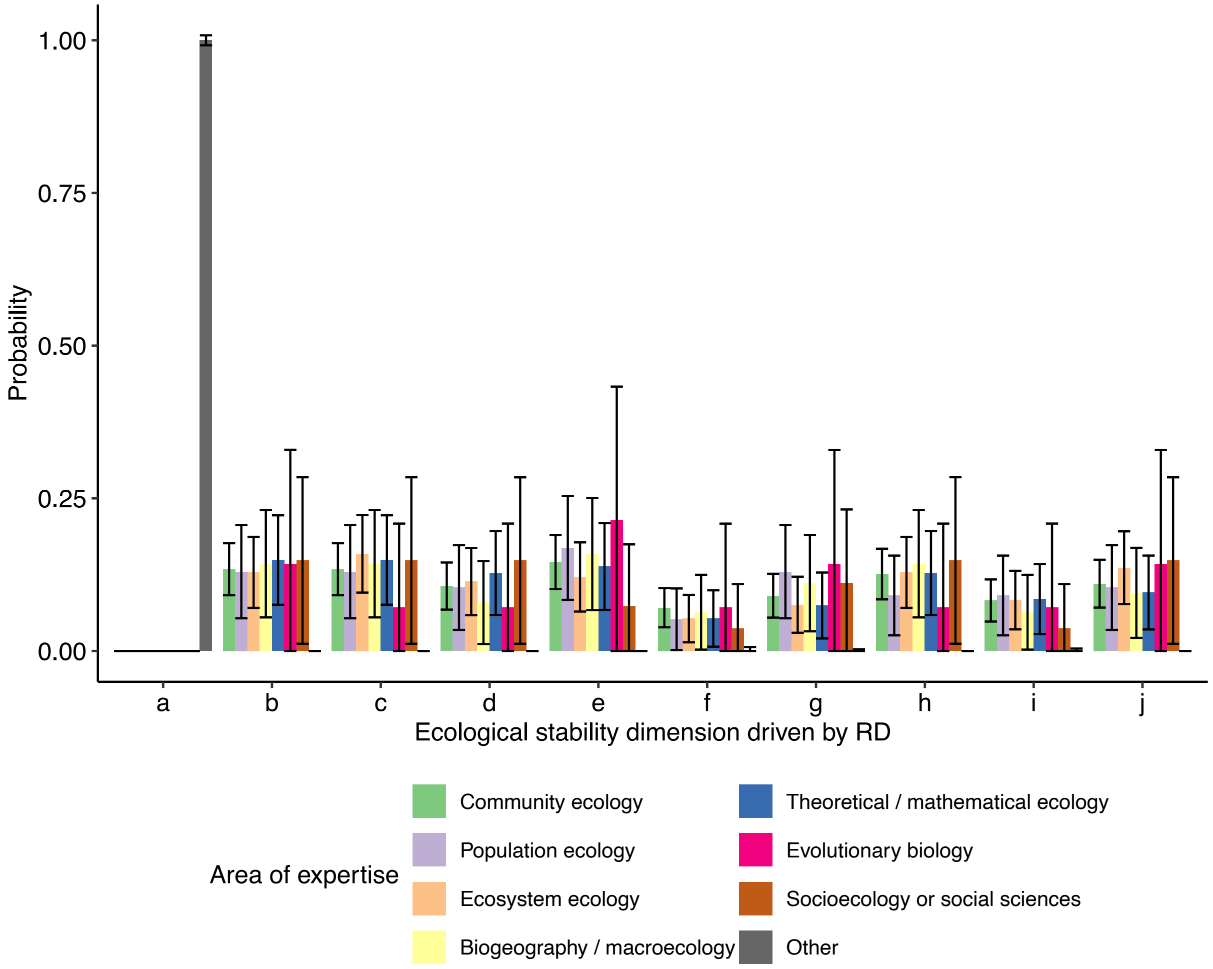


**Fig. S5.** Estimated marginal mean probabilities (bars) and 95% multinomial confidence intervals (error bars) of choosing a dimension(s) of stability driven by response diversity, RD (question 7, x-axis) and the respondents' area(s) of expertise (bar colours). Ecological stability dimensions follow Question 7 (Appendix 2), such that a = no relationship with any stability dimension; b = persistence of ecological property; c = persistence of service delivery; d = temporal variability of ecological property; e = temporal (a)synchrony; f = spatial distribution/patchiness; g = spatial (a)synchrony; h = reactivity; i = engineering resilience; and j = ecological resilience.

### Respondent's priority outcomes of Response Diversity Network

**Table S2.** Pairwise comparisons between respondent ranked preferred desired outcomes of the Response Diversity Network (Fig. 4b) using Dunn's test of multiple comparisons using Rank sums (Dinno 2024), with a Benjimini-Hochberg correction to control the experiment wise error rate. Significant ($p < 0.05$) pairwise comparisons are highlighted in Bold. Options are paraphrased here for brevity but see survey question 16 (Appendix 2) for full descriptions.

| Pairwise Comparison of Desired RDN Outcomes | Z | P.adj |
|---|---|---|
| Literature reviews- Distributed experiments | 1.724 | 0.122 |

| | | |
|---|---|---|
| Literature reviews- Grant applications | -1.517 | 0.166 |
| Distributed experiments- Grant applications | -3.240 | 0.003 |
| Literature reviews- Inclusion in monitoring | 1.551 | 0.161 |
| Distributed experiments- Inclusion in monitoring | -0.172 | 0.863 |
| Grant applications- Inclusion in monitoring | 3.068 | 0.0048 |
| Literature reviews- Inclusion in decision making | -0.793 | 0.467 |
| Distributed experiments- Inclusion in decision making | -2.517 | 0.021 |
| Grant applications- Inclusion in decision making | 0.724 | 0.497 |
| Inclusion in monitoring- Inclusion in decision making | -2.344 | 0.030 |
| Literature reviews- Data Synthesis | 4.585 | <0.001 |
| Distributed experiments- Data Synthesis | 2.861 | 0.008 |
| Grant applications- Data Synthesis | 6.102 | <0.001 |
| Inclusion in monitoring- Data Synthesis | 3.034 | 0.005 |
| Inclusion in decision making- Data Synthesis | 5.378 | <0.001 |
| Literature reviews- Other | -8.239 | <0.001 |
| Distributed experiments- Other | -9.963 | <0.001 |
| Grant applications- Other | -6.722 | <0.001 |
| Inclusion in monitoring- Other | -9.790 | <0.001 |
| Inclusion in decision making- Other | -7.446 | <0.001 |
| Data Synthesis- Other | -12.82 | <0.001 |
| Literature reviews- Methods standardisation | 0.862 | 0.437 |
| Distributed experiments- Methods standardisation | -0.861 | 0.451 |
| Grant applications- Methods standardisation | 2.379 | 0.03 |
| Inclusion in monitoring- Methods standardisation | -0.689 | 0.505 |
| Inclusion in decision making- Methods standardisation | 1.654 | 0.136 |
| Data Synthesis- Methods standardisation | -3.723 | <0.001 |
| Other- Methods standardisation | 9.100 | <0.001 |
| Literature reviews- Workshops | 2.758 | 0.010 |
| Distributed experiments- Workshops | 1.034 | 0.361 |
| Grant applications- Workshops | 4.274 | <0.001 |
| Inclusion in monitoring- Workshops | 1.207 | 0.283 |
| Inclusion in decision making- Workshops | 3.55 | <0.001 |
| Data Synthesis- Workshops | -1.827 | 0.100 |
| Other- Workshops | 11.00 | <0.001 |
| Methods standardisation- Workshops | 1.870 | 0.091 |

### Appendix 2. Survey questions from "Gauging research interest in and perceptions surrounding response diversity"

Table lists the full survey questions presented to respondents, as well as the format of each possible answer (e.g. free text, sliding scale, a single choice, choose all that apply, numeric response, etc.).

| Question | | Answer format |
|---|---|---|
| Section 1:Concepts | | |
| 1a-c | How familiar are you with work on the following topics?<br>a. Ecological stability<br>b. Functional diversity<br>c. Response diversity | Sliding scale from 1-5 as follows: 1 = Unfamiliar; 2 = Slightly familiar; 3 = Moderately familiar; 4 = Very familiar; 5 = Extremely familiar/actively work in that field |
| 2 | Do you think that response diversity is a clear concept? | Yes/No/Unsure |
| 2b | Please provide any comments related to how you think the concept should be clarified. | Free text |
| 3 | Do you think that response diversity has a widely accepted definition? | Yes/No/Unsure |
| 3b | Please give what you believe to be the widely accepted definition. | Free text |
| 4 | Which of these options (if any) captures what you consider response diversity to be in an ecological context?<br>a. Variation in how species respond to environmental change<br>b. Variation in how functionally redundant species (i.e. species contributing to the same ecosystem function(s)) respond to environmental change<br>c. Variation in how individuals respond to environmental change<br>d. Diversity of functional response traits within a functional effect group (a group of species with similar functional effect traits)<br>e. Variation in gene expression under environmental change<br>f. Other (please describe) | Choose all that apply |
| 5 | Which of these options (if any) best captures the reasons behind your interest in studying response diversity?<br>a. I am not interested in studying response diversity<br>b. To gain understanding of current response diversity levels in the community/ecosystem<br>c. To gain understanding of current ecosystem stability/resilience and the underlying mechanisms<br>d. To maximise the ability of ecosystems to cope with and/or adapt to environmental change in practice<br>e. To forecast or understand changes in ecosystem functioning under environmental change<br>f. To inform policy about important aspects of biodiversity and species to be conserved<br>g. Other (please describe) | Choose all that apply |
| 6 | Which of these options (if any) best describes how you see response diversity relating to conservation and/or ecosystem management? | Choose all that apply |

| | | |
|---|---|---|
| | a. Maintaining ecosystem functioning under environmental change by active manipulation of community composition (or genetic, or intraspecific variation)<br>b. Mitigating the impacts of environmental change on ecosystems based on predictions from response diversity theory<br>c. Monitoring response diversity as a neglected aspect of biodiversity monitoring<br>d. Other (please describe) | |
| 6b | Why? Please briefly explain what you chose this answer(s) regarding how response diversity relates to conservation and/or ecosystem management. | Free text |
| 7 | Response diversity has been proposed as a possible driver of ecological stability. Which of the following aspect(s) of stability (if any) do you think response diversity may mechanistically drive?<br>a. No relationship with ecological stability<br>b. The continuation of a community or emergent ecosystem property through time despite environmental change<br>c. The continued delivery of ecosystem services through time despite environmental change<br>d. The variability of a community or emergent ecosystem property through time<br>e. The (a)synchronicity of the temporal population dynamics of different species in a community<br>f. The distribution of species (or populations, or genes) across space (either locally or across multiple habitat patches)<br>g. The (a)synchronicity of the population dynamics of a species (or population, or gene) among different local habitat patches (connected or unconnected)<br>h. The amount of change in species' biomass or an emergent ecosystem property in response to an abrupt environmental change<br>i. The rate at which species' biomass or an emergent ecosystem property return to pre-disturbance values following an abrupt environmental change<br>j. The capacity for species' biomass or an emergent ecosystem property to return to pre-disturbance values following an abrupt environmental change or not (no return may indicate a change in ecosystem state)<br>k. Other (please describe) | Choose all that apply |
| Section 2: Studies | | |
| 8 | Have you measured and/or analysed what you would consider to be response diversity (e.g. in the field, the lab, or in simulation model scenarios)? | Yes/No/Unsure |
| 8b | If you have measured and/or analysed response diversity, then thinking about one specific study, is this work available as a preprint or journal article (with DOI)?<br>a. Yes (please provide the DOI of this study)<br>b. No | Single choice |

| | | |
|---|---|---|
| 8c | If this work is not available as a preprint or journal article, thinking about one study where you measured/analysed response diversity, in which country/countries did the study take place? | Free text |
| 8d | Thinking about this same study, please name the principal investigator(s) of this project. This data will be used to prevent double counting of responses, and will not be used for any other purpose than excluding redundant entries. | Free text |
| 8e | Thinking about this same study, in which kind of ecosystem did you primarily measure/analyse response diversity?<br>a. Marine<br>b. Freshwater<br>c. Terrestrial (tropical)<br>d. Terrestrial (temperate)<br>e. Terrestrial (dry)<br>f. Terrestrial (arctic and boreal)<br>g. Subterranean (e.g. soil)<br>h. Multiple (e.g. synthesis)<br>i. N/A (e.g. theory, sociecological analysis)<br>j. Other (please describe) | Single choice |
| 8f | Thinking about this same study, for which taxa did you primarily measure/analyse response diversity?<br>a. Plants<br>b. Microorganisms<br>c. Birds<br>d. Insects<br>e. Terrestrial mammals<br>f. Marine mammals<br>g. Fish<br>h. Reptiles<br>i. Amphibians<br>j. Multiple (e.g. synthesis)<br>k. N/A (e.g. theory, sociecological analysis)<br>l. Other (please describe) | Single choice |
| 8g | Thinking about this same study, what was the study's main objective?<br>a. To measure or estimate response diversity<br>b. To use response diversity as a predictor of another variable (e.g. ecological stability)<br>c. To use another variable (e.g. temperature, taxonomic group, etc.) to predict response diversity<br>d. Another objective involving response diversity (please describe)<br>e. The main objective did not involve response diversity<br>f. Other (please describe) | Single choice |
| 8h | Thinking about this same study, how many trophic levels did your study consider? 1 could mean within-guild, while 2 could mean predator-prey, for example. If not applicable, please write "N/A" | Free text |
| 8i | Thinking about this same study, what was the time-span of the study (in generations of your focal organism(s) or any relevant unit of time)? If not applicable, please write "N/A" | Free text |

| | | |
|---|---|---|
| 8j | Thinking about this same study, what was the spatial scale of the study (in numbers of habitat patches or any relevant unit of space)? If not applicable, please write "N/A" | Free text |
| Section 3: Future | | |
| 9 | Briefly consider an ecological community composed of a number of plant species, each of which has a slightly different preferred environmental condition. Under natural conditions, the environment (temperature, nutrient availability, weather etc.) changes on predictable cycles such as the diurnal and annual seasonal cycles, and less predictably through stochastic environmental changes to environmental conditions (e.g. an a-typically prolonged drought) and via longer-term climate change. Researchers and ecosystem managers may be interested in ensuring the continued functioning of this community and its contributions to people. In this regard they may measure, monitor, or manage for the persistence of the community through time (do populations of species persist without going locally extinct?) as well as the stability (low variability) of the total community biomass or species composition through time.<br><br>Considering the above example, at what temporal scale might you consider response diversity to be the most relevant to measure, monitor, or manage for persistence or stability in this way? | Sliding scale from 1-100, roughly: 1-20 = Very fine scales (e.g. minutes, hours); 21-40 = e.g. Daily-seasonal; 41-60 = e.g. between yrs (1-2 years); 61-80 = Broad scales (e.g. multiple years, decades, longer); 81+ = Evolutionary scales (millennia or multiple generation times) |
| 9b | How confident are you in your above answer? | Sliding scale 1-100: 1 = lowest confidence |
| 9c | Briefly, why do you consider response diversity to be most relevant at the temporal scale you chose above? | Free text |
| 10 | Considering the same example as above, at what spatial scale might you consider response diversity to be most relevant to measure, monitor, or manage for persistence or stability in this way? | Sliding scale from 1-100, roughly: 1-25 = Fine scales (e.g. microhabitats); 26-50 = Local (e.g. a single habitat patch); 51-75 = Regional (e.g. multiple connected local habitat patches); 76+ = Broad scales (e.g. multi-regional or global scale) |
| 10b | How confident are you in your above answer? | Sliding scale 1-100: 1 = lowest confidence |
| 10c | Briefly, why do you consider response diversity to be most relevant at the spatial scale you chose above? | Free text |
| 11 | Considering the same example as above, at what biological scale(s) might you consider response diversity to be most relevant to measure, monitor, or manage for persistence or stability in this way?<br>a. None of the answers capture relevant scales | Rank order. 1 = most relevant. |

| | | |
|---|---|---|
| | b. Genetic response diversity (within a population)<br>c. Metapopulation response diversity (within a species in space)<br>d. Interspecific response diversity<br>e. Response diversity within a single functional group (a group of organisms contributing to the same ecosystem function(s))<br>f. Response diversity among functional groups (groups of organisms contributing to different ecosystem functions)<br>g. Metacommunity response diversity (among communities in space)<br>h. Other (please describe) | |
| 11b | How confident are you in your above answer? | Sliding scale 1-100: 1 = lowest confidence |
| 11c | Briefly, why do you consider response diversity to be most relevant at the biological scale(s) you chose above? | Free text |
| 12 | Do you think you might measure / analyse what you consider to be response diversity in the future? | Yes/No/Unsure |
| 13 | What (if anything) do you see as the main challenge(s) facing studies of response diversity?<br>a. None of the other answers capture the main challenges<br>b. Complexity introduced by the inclusion of ecological interactions (e.g., predation, competition) in response diversity research<br>c. Confusion / lack of clarity / tautology in the definition and aims of response diversity<br>d. Interacting disturbances in natural systems (multiple stressors)<br>e. Lack of a standardised methodology for analyses<br>f. Lack of a standardised methodology for experiments<br>g. Limited data availability<br>h. Other (please describe) | Rank order: 1 = largest challenge. |
| 14 | Do you think there needs to be a standardised methodology for conducting response diversity studies? | Yes/No/Unsure |
| 14b | Why? Please explain briefly. | Free text |
| 15 | Are you aware of a standardised methodology for conducting response diversity studies? | Yes/No |
| 15b | If you are aware of a standardised methodology for conducting response diversity studies, please provide a DOI or link to an example publication using this method, or please briefly describe the method. | Free text |
| 16 | What (if anything) would you like to see as the outcome(s) of any global collaborative efforts on response diversity?<br>a. Meta-analysis / synthesis of existing data<br>b. Organised workshops (either standalone or at existing conferences)<br>c. Collaborative grant writing<br>d. Establishing a globally-distributed network of experiments (similar to e.g. NutNet)<br>e. Comprehensive literature review<br>f. Methods standardisation (e.g. establishing guidelines or development of new methods) | Rank order: 1 = most preferred outcome |

| | | |
|---|---|---|
| | g. Promoting the inclusion of response diversity in biodiversity observation and monitoring<br>h. Promoting the inclusion of response diversity in decision making and policy settings<br>i. Other (please describe) | |
| Section 4: Anonymous demographic information | | |
| 17 | What is your current career stage?<br>a. Student or recent PhD graduate (1-2 years)<br>b. 2-5 years post-PhD or equivalent experience<br>c. 5-10 years post-PhD or equivalent experience<br>d. 10+ years post-PhD or equivalent experience | Single choice |
| 18 | In which continent are you currently based? | Free text |
| 19 | Which of these options (if any) best describes your area of specialism / expertise?<br>a. Community ecology<br>b. Population ecology<br>c. Ecosystem ecology<br>d. Biogeography/macroecology<br>e. Theoretical/mathematical ecology<br>f. Evolutionary biology<br>g. Socioecology or social sciences<br>h. Other (please describe) | Choose all that apply |

### Appendix 3. Qualitative coding structure implemented for Open Ended Questions

Table presents the qualitative coding structure used by authors S.R.P-J.R. and A.L. to synthesise information from open ended questions (OEDs). See Appendix 1 for description of methods. Question numbers refer to those in Appendix 2. Codes may have sub-categories presented in brackets; for example, Q2b had higher-level codes including concept, and definition, which themselves had subcategories. Subcategories were used to extract more meaningful information than simply that the respondent was mentioning concepts/definitions (i.e. what is it about concepts/definitions they were discussing).

| Question | Number of codes | Codes |
|---|---|---|
| 2b | 5 | Concept; Definition; methods/measurement; scale; "response to what" |
| 2b (concepts) | 5 | multiple interpretations (ambiguous); missing consideration; similar to other concepts; broad concept (vague); based on unclear concepts |
| 2b (definitions) | 4 | unclear definition; multiple definitions; not linked to measurement; missing mathematical definition |
| 2b (methods) | 3 | unclear measurement; multiple measurements; not linked to definition |
| 2b (scale) | 1 | unclear scale |
| 3b | 4 | definition; organism; species; community; functional group |
| 4 (Other) | 4 | "community"; "emergent property"; "and other species"; all have been used |
| 6 (Other) | 5 | characterizing disturbance; actively manage for high response diversity; guide conversations; understanding resilience; conservation prioritization |
| 6b | 5 | lack of practical applications; skepticism/limitations of active manipulation; belief that RD is a useful indicator to understand and manage change; To understand whether RD is a useful indicator to understand and manage change; all options important |
| 7 (Other) | 3 | temporal; not a driver; environmental variability |
| 9c | 7 | selected based on characteristics of the community; selected based on characteristics of the environment/disturbance; selected based on human-centered outcomes; selected based on practical/logistical considerations; selected to account for evolutionary processes; selected to account for stochasticity; mentions duration (in addition to frequency) |

| | | |
|---|---|---|
| 10c | 5 | selected based on characteristics of the community; selected based on characteristics of the environment/disturbance; selected based on human-centered outcomes; selected based on practical/logistical considerations; discusses multiple scales/cross-scales |
| 11c | 5 | selected based on relevance to EF, stability, or resilience; selected based on characteristics of the community; selected based on human-centered outcomes; selected based on practical/logistical considerations; discusses multiple scales/cross-scales |
| 13 (Other) | 2 | methodological challenges; conceptual challenges |
| 14b | 5 | need for greater comparability/standardization; one method is difficult/impractical; one method is constraining/reduces plurality; one method cannot account for different systems; one method is not needed (generic) |
